# Convergence of immune escape strategies highlights plasticity of SARS-CoV-2 spike

**DOI:** 10.1101/2022.03.31.486561

**Authors:** Xiaodi Yu, Jarek Juraszek, Lucy Rutten, Mark J. G. Bakkers, Sven Blokland, Niels J.F. van den Broek, Annemiek Y.W. Verwilligen, Pravien Abeywickrema, Johan Vingerhoets, Jean-Marc Neefs, Shah A. Mohamed Bakhash, Pavitra Roychoudhury, Alex Greninger, Sujata Sharma, Johannes P. M. Langedijk

## Abstract

The SARS-CoV-2 spike protein is the target of neutralizing antibodies and the immunogen used in all currently approved vaccines. The global spread of the virus has resulted in emergence of lineages which are of concern for the effectiveness of immunotherapies and vaccines based on the early Wuhan isolate. Here we describe two SARS-CoV-2 isolates with large deletions in the N-terminal domain (NTD) of the spike. Cryo-EM structural analysis showed that the deletions result in complete reshaping of the antigenic surface of the NTD supersite. The remodeling of the NTD affects binding of all tested NTD-specific antibodies in and outside of the NTD supersite for both spike variants. A unique escape mechanism with high antigenic impact observed in the ΔN135 variant was based on the loss of the Cys15-Cys136 disulfide due to the P9L-mediated shift of the signal peptide cleavage site and deletion of residues 136-144. Although the observed large loop and disulfide deletions are rare, similar modifications became independently established in several other lineages, highlighting the possibility of a general escape mechanism via the NTD supersite. The observed plasticity of the NTD foreshadows its broad potential for immune escape with the continued spread of SARS-CoV-2.

## Introduction

The viral surface spike (S) protein of SARS-CoV-2 is critical for the viral life cycle, the primary target of neutralizing antibodies (*1–4*) and a key target for prophylactic vaccines. S is a large, trimeric glycoprotein that mediates both binding to host cell receptors and fusion of the viral and host cell membranes through its S1 and S2 subunits respectively (*5–7*). The S1 subunit comprises two distinct domains: an N-terminal domain (NTD) and a host cell receptor-binding domain (RBD) which are both targets of neutralizing antibodies and escape mutations are described for both regions (*8*). The immunodominant NTD binds antibodies with high neutralizing and protective potential (*2, 9–15*) and most SARS-CoV-2 variants have small deletions in the exposed protruding loops of NTD (*16–19*). In this study we characterize spikes of two isolates, obtained from samples from infected individuals in Peru (ΔN25) and Brazil (ΔN135) in January 2021, both containing large deletions in the NTD. Additionally, the ΔN135 isolate contains mutations in the RBD and a mutation in the signal peptide that together with the deletions result in a major remodeling of the structure of the NTD due to loss of the 15-136 disulfide (DS_15-136_). Both S proteins fold correctly and maintain fusion capacity despite the disulfide loss and large deletions in a small beta-sheet on top of the NTD galectin fold (β_N3N5_). High resolution single-particle electron cryo-microscopy (Cryo-EM) structures supplemented with antigenicity profiling underline the potential impact of these deletions on immune escape.

## Results

Next-generation sequencing analysis of SARS-CoV-2 RNA isolated from nasal swab samples collected from study participants in the Phase 3 trial of the Ad26.COV2.S vaccine (VAC31518COV3001, Ensemble, funded by Janssen Research and Development and others, ClinicalTrials.gov number NCT04505722(*20*)) revealed various adaptations in the S gene sequences. Multiple study participants from Peru and one from Argentina showed common mutations in the NTD and the RBD and a unique large deletion of residues 63-75 in the N2 loop of the spike. Since the spike has the deletion in the N5 loop common for C37 and the novel N2 loop deletion, the spike is named ΔN25 (Fig 1 and Table S1). Samples obtained from two study participants that were taken on January 12^th^ and 17^th^ of 2021 in Sao Paolo, Brazil, showed identical amino acid sequences for the S protein that were very different from the global consensus. Apart from several earlier described mutations in the RBD, these sequences showed a mutation in the signal peptide and two large deletions in the NTD of residues 136-144, a beta-strand preceding the N3 loop, and residues 258-264 in the N5-loop and therefore this spike is named ΔN135 (Fig. 1, Table S1).

**Figure 1.**
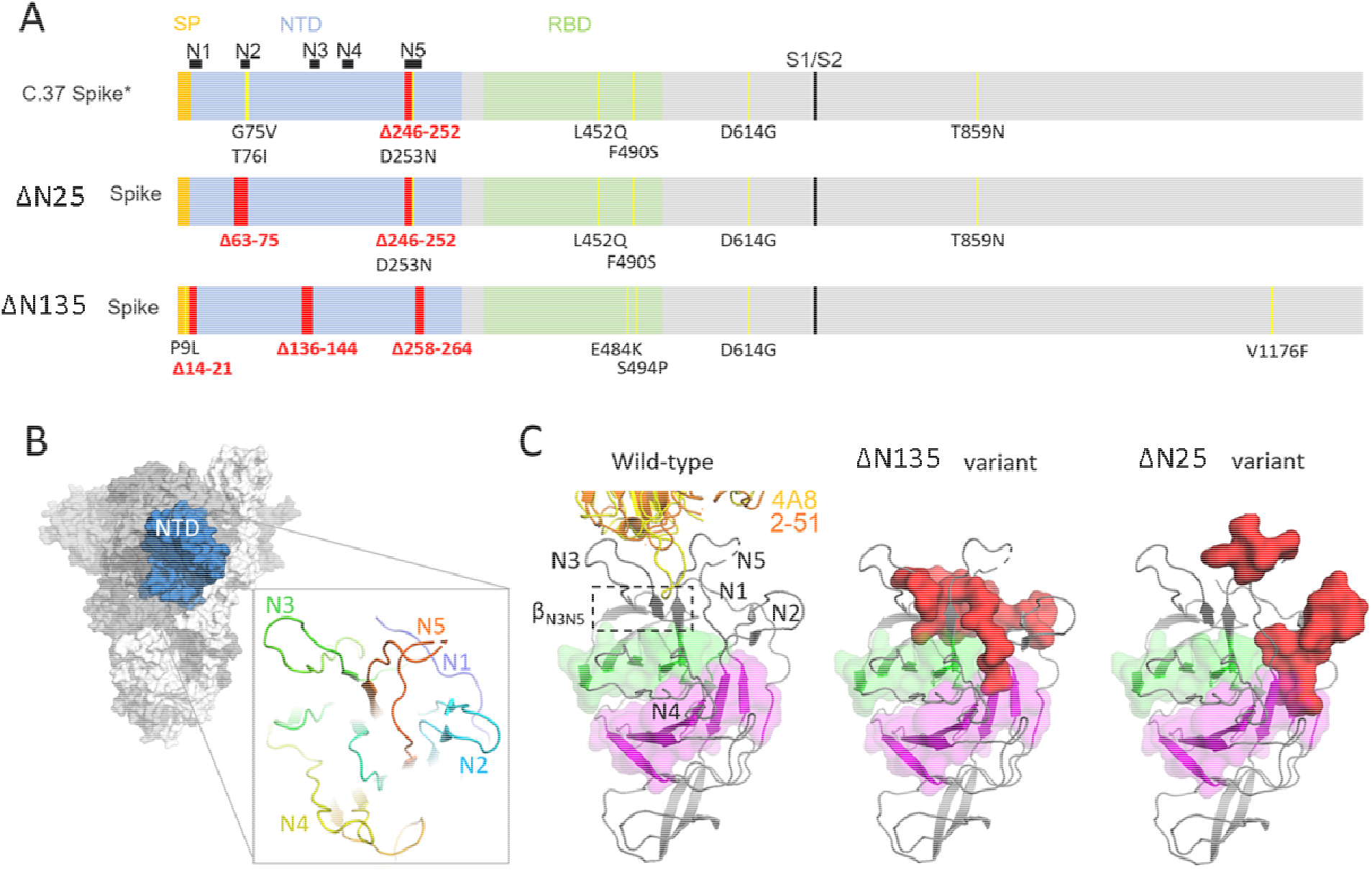
**a** Schematic representation of the C.37, ΔN25 and ΔN135 spikes with signal peptide (SP) indicated in yellow, N-terminal domain (NTD) in blue with the five NTD loops indicated above the bar, receptor binding domain (RBD) in green, S1/S2 cleavage site in black and mutations in yellow and deletions in red. **b** Sideview of a spike with the NTD domain in blue. NTD loops N1 (blue), N2 (cyan), N3 (green), N4 (yellow) and N5 (orange) are plotted in the inset as ribbon **c** Left panel: sideview of the NTD with the two sheets of the galectin-fold in green and magenta and indicated N-loops. The β_N3N5_ sheet on top of the galectin-fold is boxed with a dashed line. As a reference for the NTD supersite, structures of Fabs 4A8(*10*) (PDBID 7C2L) ^1^ and 2-51 (PDBID 7L2C) (*14*) are indicated in yellow and orange ribbons. In the middle and right panels, the deleted amino acids are depicted for the ΔN25 and ΔN135 spikes in red as spacefilling representation.

### Variant spikes remain fusogenic

Given the extensive changes that ΔN25 and ΔN135 spikes had accumulated compared to the original SARS-CoV-2 strain, we attempted to confirm their ability to successfully accomplish membrane fusion. We measured the impact of the changes in the full-length variant spikes on fusion activity compared with the wild-type Wuhan-Hu-1 (GenBank accession number: MN908947) in a cell-cell fusion assay that makes use of a fluorescent reporter protein to visualize syncytia formation (*21*). HEK293 cells were transiently transfected with plasmids encoding S, ACE2, TMPRSS2 and GFP. Transfection of GFP alone, or of a prefusion-stabilized S protein did not yield syncytia. On the contrary, major syncytia formation was observed with the Wuhan-Hu-1 S protein. Likewise, when cells were transfected with either one of the two variant S proteins, clear syncytia were visible. These data demonstrate that the variant S proteins remain fully functional despite considerable changes in the NTD (Fig 2.A).

**Figure 2.**
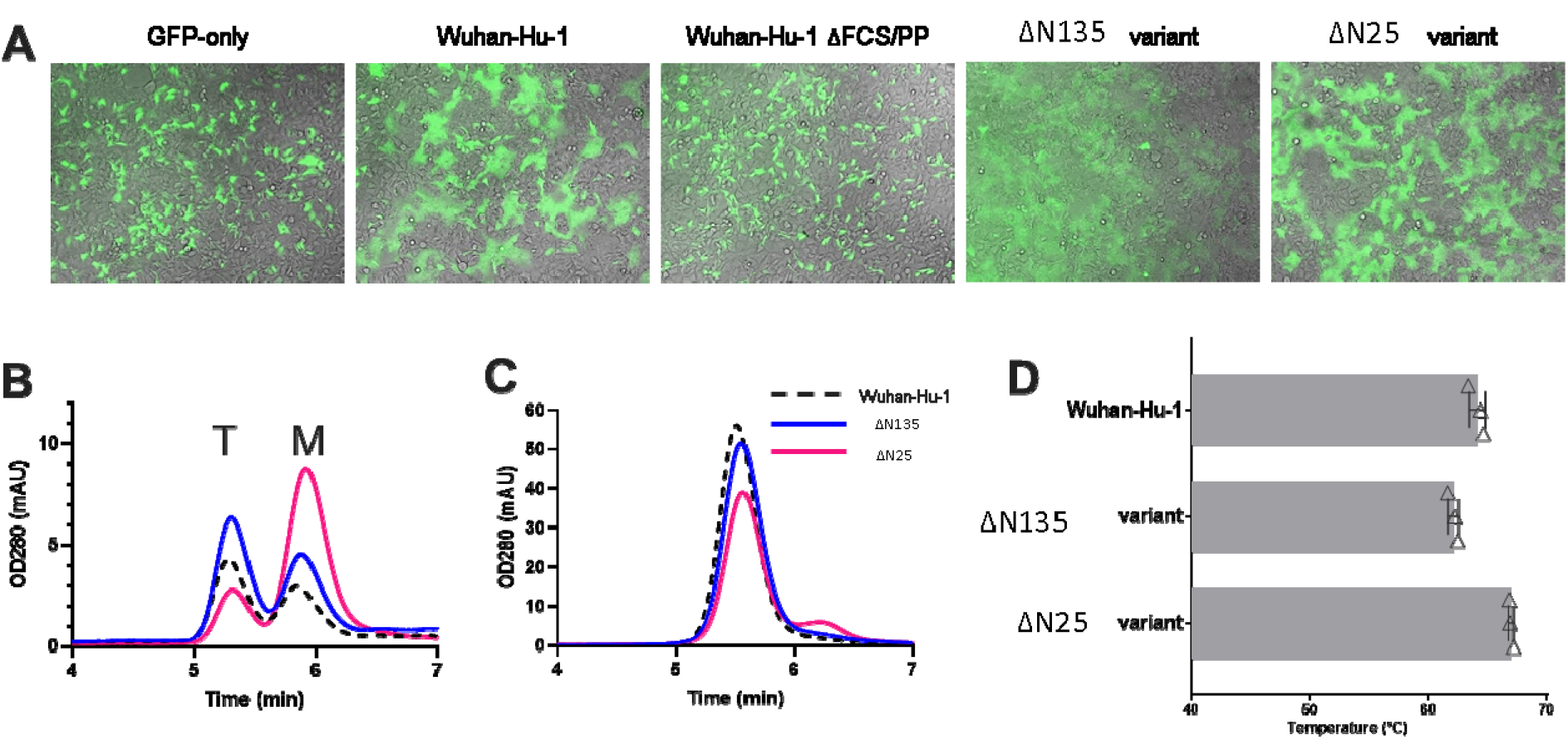
Characterization of variant spikes. **a** Cell-cell fusion assay in HEK293 cells by co-transfection of plasmids encoding S protein, ACE2, TMPRSS2, and GFP. Shown are overlays of the GFP and brightfield channels 24□hr after transfection. The different S protein constructs are indicated; ‘GFP-only’ did not include S plasmid. (b) Analytical SEC chromatograms of the S Wuhan-Hu-1 S variant (black dotted line), the ΔN135 S variant (blue line) and the ΔN25 S variant (magenta line) in cell culture supernatants on an SRT-10C SEC-500 15 cm column. The T indicates the trimer peak and the M indicates the monomer peak. (c) SEC chromatograms of the purified trimers of the S variants. Coloring is the same as in figure 2b. (d) Main melting event temperatures (TM_50_) of the S protein variants. Data are represented as mean + SD of n = 3 replicates.

### Characterization of the ΔN25 and ΔN135 spikes

We designed soluble versions of the variant S proteins and produced them in transiently transfected expi293F cells to enable biochemical and structural characterization. To obtain high quality S proteins with reasonable yields, the furin cleavage site was mutated and stabilizing substitutions to proline were added at positions 892, 987, and 942 in the S2 domain(*22*). The variant spikes were produced at levels comparable to the Wuhan spike in the crude cell culture supernatant (Fig 2B). The quaternary structure of the ΔN25 spike was less stable and showed a higher fraction of monomeric S compared to the ΔN135 and Wuhan variants. After purification, only trimeric S proteins remained (Fig 2C). These purified proteins were used for all subsequent experiments. All three S proteins showed the typical minor melting event at approximately 49°C and a higher main melting event that differed among the spikes. The *Tm*_50_ of the ΔN25 spike was 2.5°C higher, and that of the ΔN135 spike was 2.5°C lower, as compared to the Wuhan spike (Fig 2D, Supplementary Figure 1).

### Antigenicity of the variant spikes

To investigate the impact of the variant point mutations and deletions on the antigenicity, we measured binding of a selection of MAbs to the ΔN25 and ΔN135 spikes and compared it with the binding to the Wuhan-Hu-1 spike. The antigenic assessment was performed using biolayer interferometry to measure S protein binding to ACE2-Fc and a panel of six SARS-CoV-2 neutralizing antibodies directed against the RBD (S2M11, S2E12, C144, 2-43, S309 and COVA2-15 (*2, 23–26*)), three neutralizing antibodies against the supersite of the NTD (2-51, COVA1-22 and 4A8 (*10, 14, 26*)) and a non-neutralizing antibody against the lower part of the NTD (DH1055(*11*)) (Fig. 3). The NTD-specific antibodies lost all binding to both variant spikes, except for some residual binding of DH1055 to the ΔN135 S protein. Although ACE2-Fc was still able to bind, MAbs 2-43 and COVA2-15 lost all binding to the variant spikes. Binding to the RBD of the ΔN135 variant was most significantly impacted and out of the entire panel, only S2E12 and S309 antibodies directed against conserved RBD sites were not or hardly affected. The loss of binding to the RBD is most likely caused by the E484K mutation, which is part of the epitopes of the MAbs SM11, 2-43, C144 and COVA2-15 (*27–29*).

**Figure 3.**
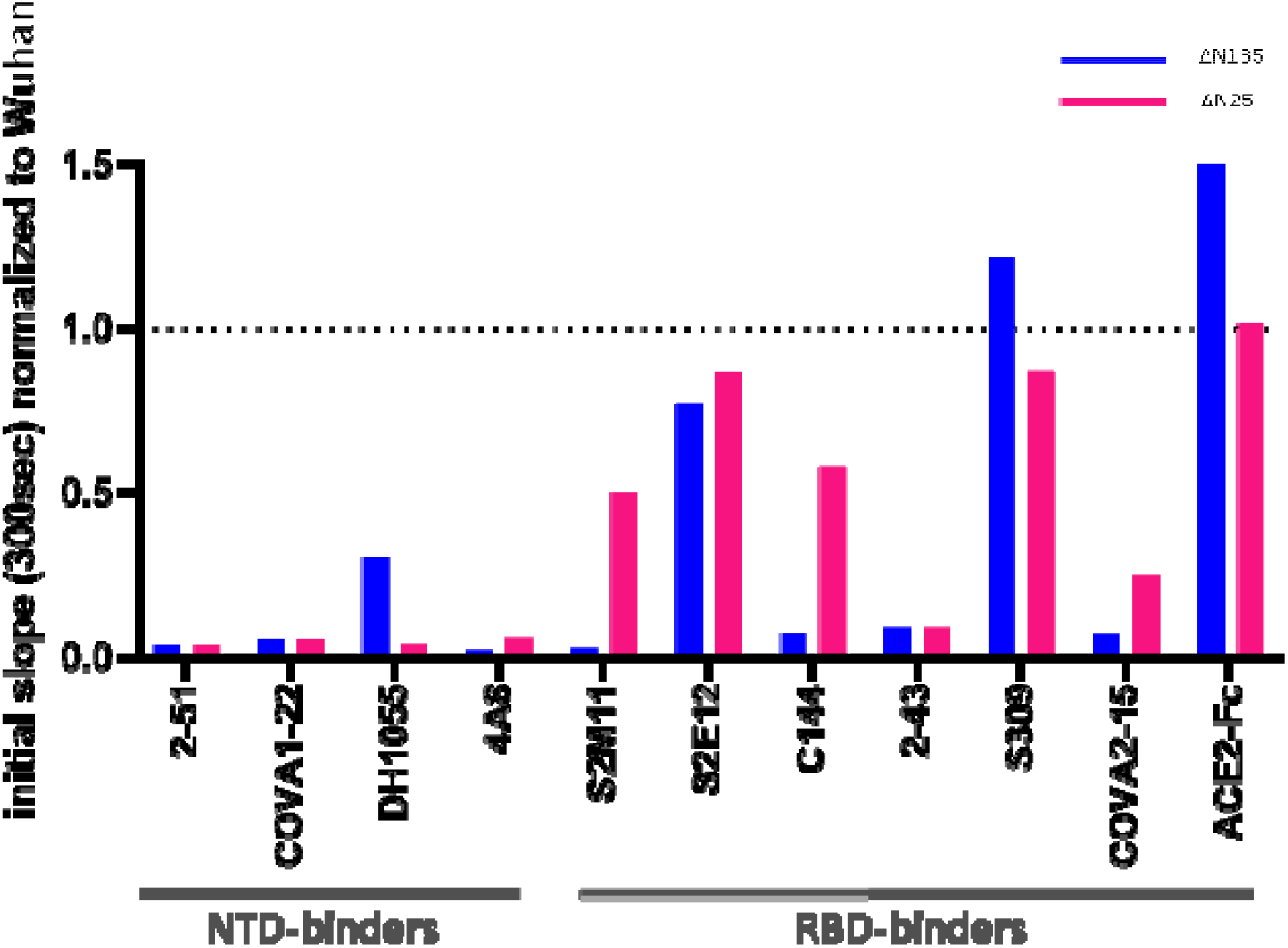
Biolayer Interferometry. Binding of NTD and RBD-specific MAbs and ACE2-Fc to purified ΔN135 (blue) and ΔN25 (magenta) S trimer variants measured with BioLayer Interferometry, showing the initial slope V0 at the start of binding, normalized to that of the Wuhan variant (dashed line).

### Shift in signal peptide cleavage site and subsequent loss of disulfide

In SARS-CoV-2 S, a conserved cysteine Cys15 is present near the N-terminus of the mature protein and forms a disulfide bond with Cys136. Only in the case of the branch of coronaviruses that includes SARS-CoV-2 S, the cysteine is located almost directly adjacent, two amino-acids away from the signal peptide (SP) cleavage site (red in Fig. S2). Mutations in the signal peptide that shift the cleavage position downstream of Cys15, would prevent disulfide DS_15-136_ from forming and consequently impact the structural architecture of the NTD. To investigate the effect of the signal peptide P9L mutation, we performed liquid chromatography-mass spectrometry (LC-MS/MS) from a tryptic digest of purified Wuhan-Hu-1 and the ΔN135 S protein to determine the N-terminal residue of the mature proteins. We found that, in line with published observations(*21*), the Wuhan-Hu-1 S protein was cleaved after position 13 (Fig. 4a, Fig. S3). In contrast, for the ΔN135 S protein no peptides were detected up to N-terminal residue 22. Whereas the SignalP-6.0 prediction software predicted the loss of the cysteine by cleavage directly C-terminal to Cys15, according to LC-MS/MS the N-terminus is truncated by 7 additional residues (Fig. 4a). Interestingly, in the ΔN135 spike the loss of Cys15 is accompanied by the loss of Cys136 due to the large deletion of residues 136-144. The loss of both cysteines could indicate a compensatory mutation since an unpaired cysteine can impact correct folding of the spike.

**Figure 4.**
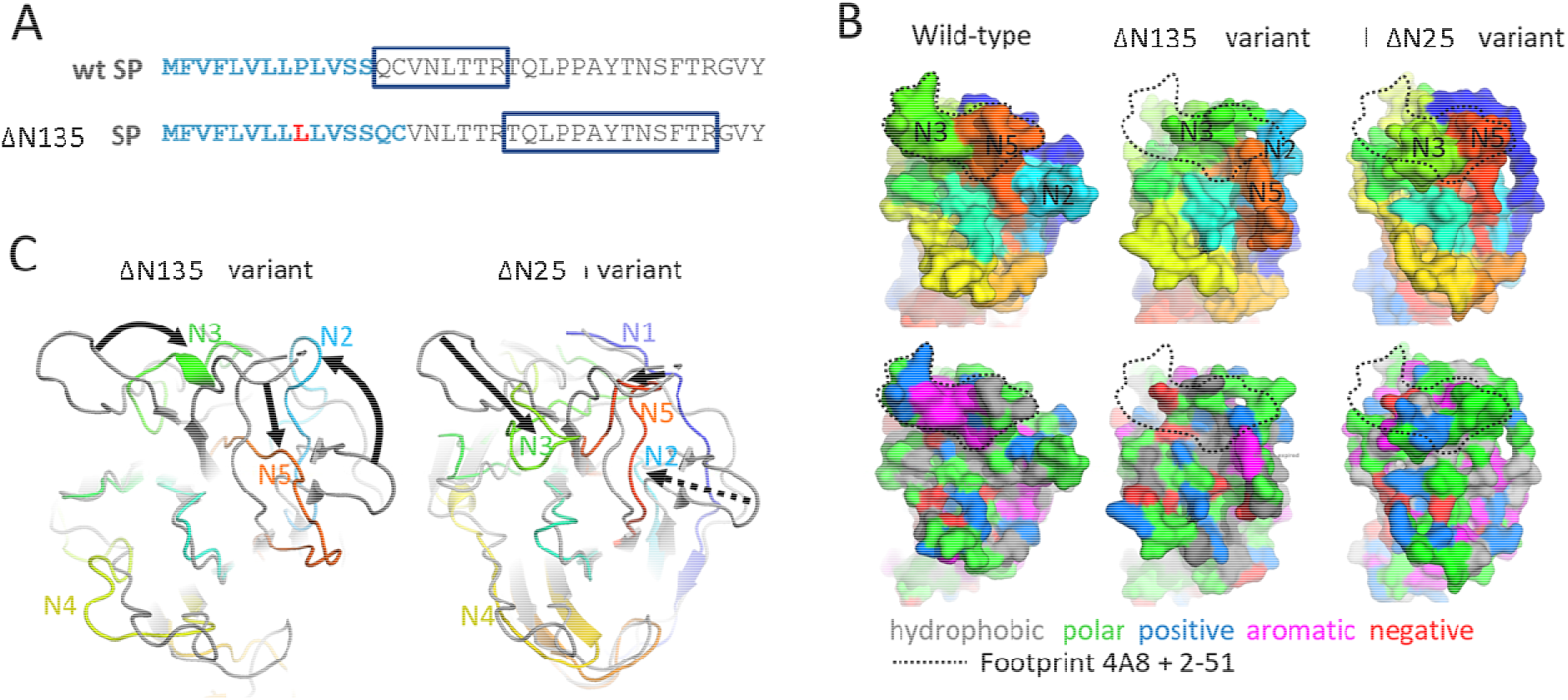
Conformational plasticity of NTD in spike variants. **A.** The signal peptide predicted by SignalP-6.0 of the wt SP and the ΔN135 SP are shown in blue bold characters. The mutation in the ΔN135 S is shown in red. Most N-terminal peptide detected using mass spectrometry is boxed. The peptides QCVNLTTR, VNLTTR or NLTTR are not found for the ΔN135variant S. B. Surface representation of the NTD supersite (same view as Fig. 1B). The color code in the upper panels is the same as in Fig. 1B. The color code in the bottom panels is based on residue type – hydrophobic in grey, polar in green, positively charged in blue, negatively charged in red and aromatic in magenta. Dashed contours indicate the joint footprint of MAbs 4A8 plus 2-51 on the reference spike. The epitope contour was also plotted over the variant NTDs as guidance to indicate the changes introduced by the deletions. C. Superposition of ribbon representation of the reference NTD (grey, PDBID 7C2L) and variant NTDs with colors and view as in Fig 1B. Arrows indicate rearrangement of the loops as a result of the deletions. The dashed arrow indicates the complete deletion of the N2 loop in the ΔN25 variant.

### CryoEM analysis of variant spikes

To understand the structural impact of the large NTD deletions and the loss of DS_15-136_ (ΔDS_15-136_) in the Brazilian variant, we solved the structures of the stabilized ectodomains of both spike variants by CryoEM analysis. The overall structure of the ΔN25 and ΔN135 trimers are the same as that of the Wuhan spike with the D614G mutation except for the loops in the NTD (Fig. 4 bc, Fig S4). From the ΔN25 spike dataset, one stable class with one RBD-up was able to be refined into high resolution (Table S2, Fig S5a, S6). The ΔN25 spike has a 7-residue deletion in the N5 loop typical for the C.37 lineage(*30*). As a result of this deletion and the complete loss of the N2-loop due to the large 13-residue deletion of residues 63-75, the N5-loop shifts towards the N2 and N1 loops and concomitantly, the N3-loop shifts to a position previously occupied by N5. As a result of the deletions and N-loop shifts, the 3-strand β-sheet formed by N3 hairpin and N5 (β_N3N5_) on top of the galectin-fold is lost and as a result, the N4-loop is shifted away from the other loops. The deletions and remodeling of N2, N3, N4 and N5 result in major antigenic changes in the NTD supersite (Fig. 4 b and c). Compared with the Wuhan spike with the same stabilizing mutations, the ΔN135 variant is more open. It acquires predominantly the 1-RBD up conformation (73% 1-up, 23% down) compared to 20% 1-up, 80% down for the Wuhan variant (Table S2, Fig. S5b, S6). This increase in the RBD ‘up’ state is likely due to the E484K mutation, previously described to influence this balance (*31*). Deletion of N1 results in loss of DS_15-136_ and exposes a hydrophobic patch which contributes to a large reorganization of the NTD loops. The conserved N2 loop has completely shifted position and occupies the space of the deleted N1 loop (Fig 4c). The deletion of one of the strands of the N3 beta-hairpin destroys the 3-strand β-sheet β_N3N5_ (Fig.1C). As a result, N3 completely shifts and occupies the space of the deleted N1 loop. Finally, the deletion in N5 and the loss of the secondary structure of β_N3N5_ results in a shift of N5 to the space previously occupied by N2 and N4 shifts away from the other loops. The loss of DS_15-136_ and β_N3N5_ due to the deletions in N1, N3 and N5 causes a dramatic remodeling of the N2, N3, N4 and N5 loops that includee the NTD supersite (Fig. 4bc) and a reduced stability of the spike (Fig. 2D).

### Spread of the DS_15-136_ breaking mutations

P9L and the previously described S13I (*32*) cause a shift in signal peptide cleavage, resulting in the loss of Cys15. This SP-shift can be indirectly detected by the loss of binding to MAb COVA1-22 (Fig. 5) which depends on the NTD N-terminus. A panel of common SP mutations, including P9L and S13I, was evaluated for Mab COVA1-22 binding to investigate the occurrence of both the signal peptide cleavage shift and concomitant loss of DS_15-136_. Apart from P9L, S13I and C15F, only S12P resulted in reduced COVA1-22 binding which agreed with the predicted signal peptide cleavage shift and concomitant loss of Cys15 according to the SignalP - 6.0 software (*33*) (Figure 5).

**Figure 5.**
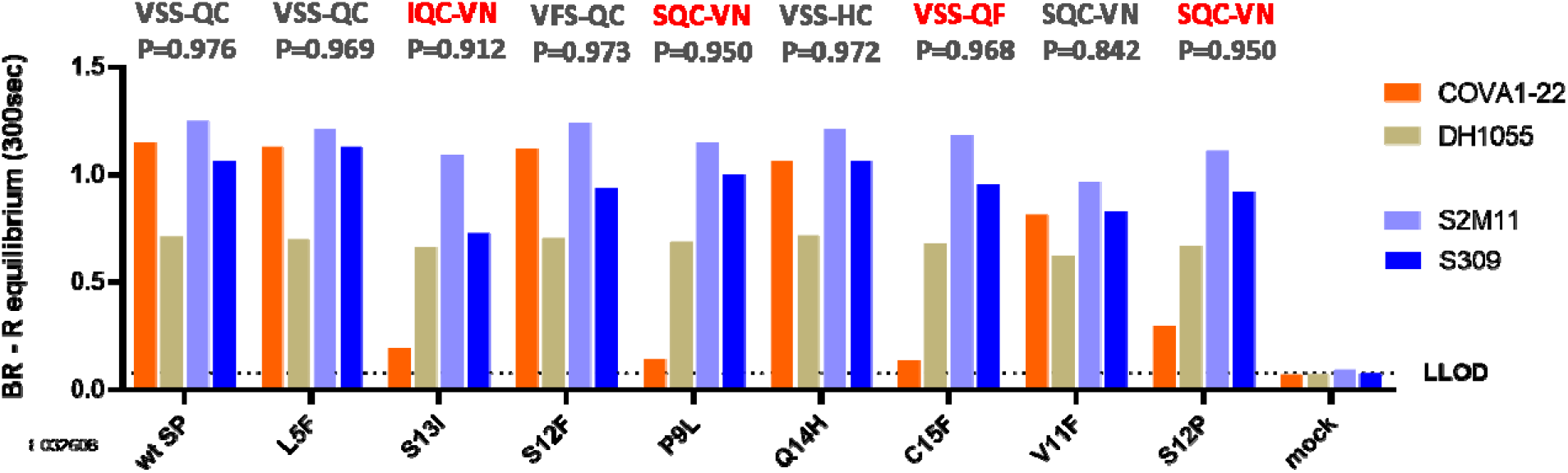
Impact of SP mutations on Spike NTD antigenicity. Binding of Mabs COVA1-22, DH1055, S2M11 and S309 to the S trimer with the wild type signal peptide (wt SP) and with different mutations in or just after the signal peptide, measured with Biolayer Interferometry using Octet. The R equilibrium calculated at 300 seconds is shown as bars. Probability (P) of the signal peptide cleavage site as predicted by SignalP 6.0 is shown above the bar graph. Mutations that result in loss of Cys15-Cys136 are indicated in red. Octet analysis was performed on crude cell culture supernatants.

NTD is a hotspot for deletions in the S protein, and the same deletions keep evolving on independent branches of the phylogenetic tree of S (Fig 6A). ΔDS_15-136_ can occur via mutation or deletion of either of the two cysteine residues (Fig. 6B). S13I and P9L are the most frequent causes for the loss of Cys15 via the cleavage site shift mechanism, but direct mutation of Cys15 is also observed (Supplementary Table 3). Cys136 is removed only via direct mutation and occurs less often. Approximately half of the lineages with ΔDS_15-136_ have both cysteines removed as in the Russian AT.1 lineage (*34*) or the C1.2 lineage (*35*). The distribution of the ΔDS_15-136_ variants on the phylogenetic tree of SARS-CoV-2 S (Fig. 6C) and the different paths leading to the disulfide loss (Table S3) suggest that ΔDS_15-136_ could have evolved in multiple lineages independently, and in several cases became dominant within the lineages. Figure 6d shows the most significant incidences of ΔDS_15-136_ in SARS-CoV-2 lineages. Before the Delta became dominant and outcompeted many of these lineages, in many cases, percentage of ΔDS_15-136_ showed an ascending trend. After replacement of most of the strains by Delta and subsequently Omicron lineages, once again, ΔDS_15-136_ is reemerging in diverse geographical locations (Table S4).

**Figure 6.**
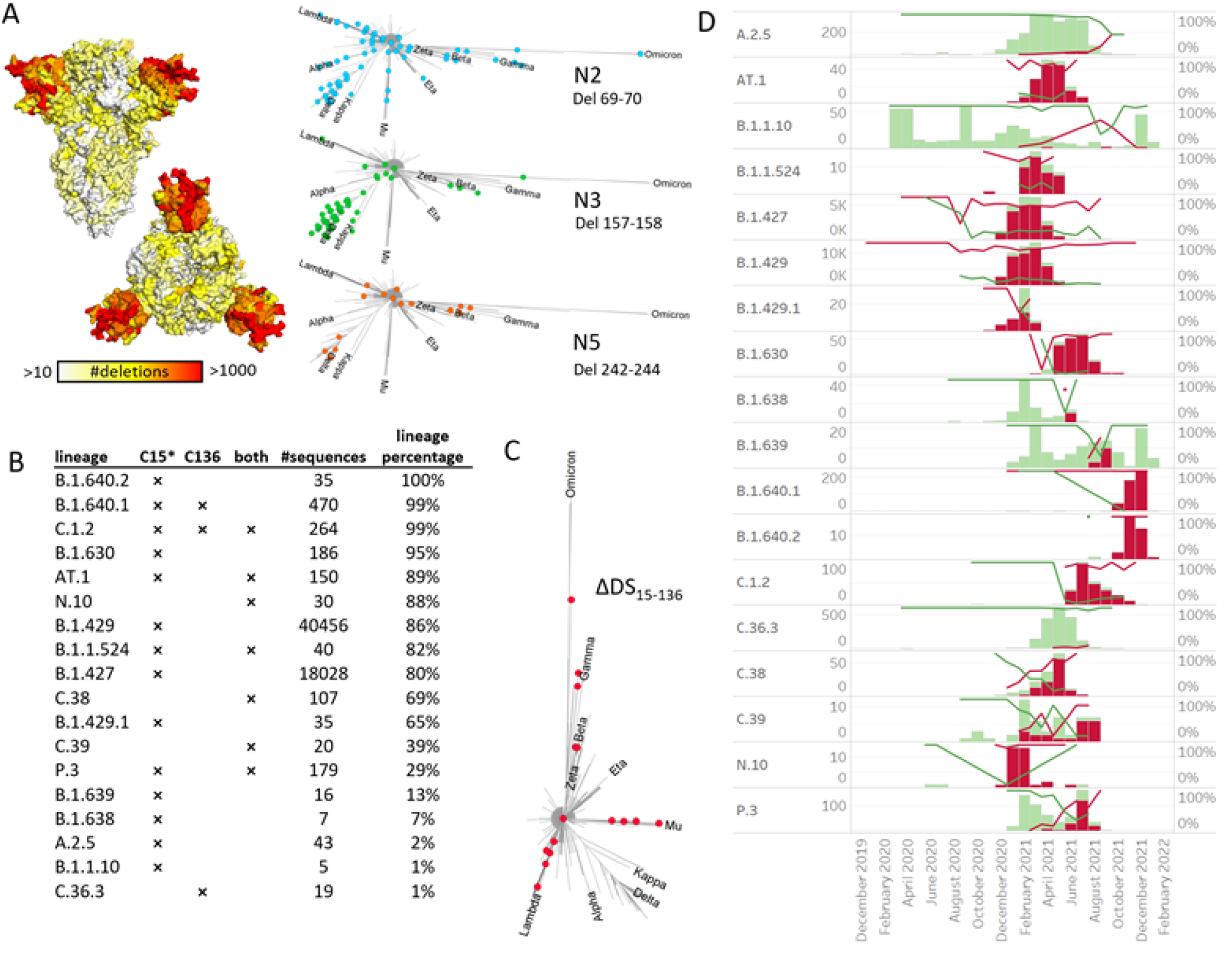
NTD deletions and occurrences of DS_15-136_ loss in lineages. A. Deletion frequency based on GISAID 25 Jan 2022, plotted on spike surface (side and top view). White corresponds to positions with less than 10 deletions registered on GISAID, while red is assigned to positions with more than 1000 deletions observed. In the right panels, deletions in N2, N3 and N5 loops are plotted on the S-protein phylogenetic trees (see Materials and Methods section), with the major variants of concern indicated as reference. Only deletions that were identified in more than 1% of their respective lineage sequences are plotted. B. List of lineages containing at least 1% of ΔDS_15-136_ variants. For each lineage, the mechanism of the DS_15-136_ loss is indicated with a cross, where C15* stands for both a direct C15 mutation or deletion, or signal peptide mediated cleavage site shift. C136 stands for C136 mutation, and “both” indicates both cysteines were removed via any of the possible mechanisms. The fraction of ΔDS_15-136_ sequences within each lineage was calculated in the last column. C. Lineages containing at least 1% of ΔDS_15-136_ variants plotted on the phylogenetic tree of SARS-CoV-2 S. D. Time evolution of the ΔDS_15-136_ containing lineages, with variants containing the cystine bridge colored in green and variants with the cystine bridge absent colored in red. The bars depict lineage counts for each month (left axis), and the lines – percentages within each lineage of both sub-variants (right axis).

## Discussion

The rapid global spread of SARS-CoV-2 leads to recurrent emergence of variants with either higher transmissibility or decreased recognition by protective immune response. The NTD undergoes rapid antigenic drift and accumulates a larger number of mutations and especially deletions relative to other regions of the spike (Fig 6A). In this study, we describe two spike variants, one from Peru and one from Brazil with typical point mutations in the RBD but dramatic and rare deletions in the NTD (Fig 1). Since the observed deletions are extensive, we examined folding and function of the variant spikes and investigated their structural impact. Both spikes showed robust expression and maintained fusogenicity, and the purified soluble proteins showed comparable thermostability and ACE2 binding (Fig 2, Fig S1). As a result of deletions, both spikes show complete loss of antibody binding to the NTD supersite (Fig 3, Fig 4). Additionally, the mutations in the ΔN135 spike impacted binding of most of the RBD specific antibodies (Fig 3). The ΔN25 variant derived from the C.37 lineage, a variant of concern (VOC) with a large 7-residue deletion in the N5 loop (*30*) acquired an additional 13-residue deletion in the N2 loop compared to C37. The ΔN135 variant belonging to the B.1.1.294 lineage acquired three large deletions: a 9- and a 7-residue deletion in the N3 and N5 loop respectively, and a deletion of the N-terminus as a result of signal peptide cleavage shift leading to the DS_15-136_ loss. Structural analysis of the proteins using CryoEM showed that the overall fold of the spikes was maintained and the galectin-fold of the NTD remained intact despite the large deletions and loss of the disulfide bridge (Fig S4, Fig S5). However, the loops that constitute the NTD supersite were completely remodeled or relocated in both proteins (Fig 4), which explains the dramatic changes to the NTD antigenicity profile. In the ΔN25 spike complete deletion of the N2 and partial deletion of N5 loop results in large shift of the N3 and N4 loops. In the ΔN135 spike, N2 and N3 move to the position of the deleted N1 and N4 moves away from the other loops. The relocation of the loops was enabled by the loss of the β_N_3N_5_ β-sheet due to deletion of the N3 β- hairpin and the deletion in the N5 loop.

Aside of the extensive loop deletions, the virus can remodel the NTD supersite by shifting its signal peptide cleavage site with the P9L point mutation. We experimentally verified that the mutation causes a longer truncation of the N-terminus by Mass spectrometry of tryptic digests, loss of binding to MAb COVA1-22 specific for the NTD N-terminus and by the CryoEM structure determination (Figs 3, 4, 5). S13I and to a lesser extend S12P also cause the peptide cleavage shift (Fig. 5) (*32*). Next to the direct mutation or deletion of one of the cysteines, the signal peptide mutations constitute an additional mechanism via which ΔDS_15-136_ can occur.

The mutations that shift the cleavage site, together with the Cys15 and Cys136 mutations and deletions were used to identify ΔDS_15-136_ variants in the GISAID database (Supplementary Table S3, Fig 6B, Fig 6C). Although these modifications are relatively rare, ΔDS_15-136_ is widespread both geographically and in terms of occurrences on the phylogenetic tree of S. This new escape mechanism arose independently in different geographical locations and even became dominant in some lineages until Delta replaced most other variants around the world. However recently, in the midst of the ongoing Omicron wave, Colson et al (*36*) reported an emergence of a new concerning variant (B.1.640.2) in Southern France, probably of Cameroonian origin which also evolved the ΔDS_15-136_ feature.

In the last two years, the NTD domain of the SARS-CoV-2 spike has been confirmed as a hotspot for deletions (Fig 6A). Within NTD, deletions are further clustered around a few sites: residues 69-70, 141-143, 156-159 and 242-245. Deletions at these sites recur independently in large number of unrelated lineages, as depicted in the phylogenetic trees of SARS-CoV-2 S in Fig 6A. The large capacity for deletions in N2, N3 and N5 loops together with the ability to remove N1 with the ΔDS_15-136_ mechanism to further rearrange all surrounding loops allows the virus to completely remodel the NTD supersite, as depicted in Fig 4 and Fig. S7. Moreover, the mechanism of reshaping the loops via ΔDS_15-136_ seems to have evolved independently in multiple branches of the SARS-CoV-2 phylogenetic tree, suggesting this important escape mechanism may also play a role in the future variants of concern.

As collective immunity to the virus grows, immune evasion will likely become an important fitness advantage, as recently observed for the Omicron variant. It is likely that escapes via structurally tolerated large deletions and/or the ΔDS_15-136_ mechanism will occur again when selection based on immune evasion continues. In fact, deletions of the loops are already firmly incorporated in the Delta and Omicron lineages. ΔDS_15-136_ has also been registered in these variants of concern albeit at low frequencies. When analyzed locally (Supplementary Table S4), at the end of the Delta wave Delta lineages in Sweden and Chile started to develop ΔDS_15-136_. With the rise of Omicron these lineages were eventually outcompeted, but the first cases of Omicron BA.1 and BA.1.1 ΔDS_15-136_ have also recently been registered in some US states. With increasing global immunity, the escape mechanisms that are currently rare, should be closely monitored and it would be important to understand the constraints of the NTD erosion and the balance between NTD function and structural integrity.

## Acknowledgements

We thank Lam Le and Pascale Boucher for technical support. We would like to thank Marit van Gils for kindly providing COVA1-22 and COVA2-15.

## Author contribution

X.Y., J.J., L.R., M.J.G.B. and J.P.L designed the study, X.Y., J.J., L.R., M.J.G.B, S.B., N.J.F.vdB, A.Y.W.V., P.A., J.V., J.N., planned and / or performed biochemical assays and purifications, X.Y. and P.A performed EM sample preparation, data collection, data processing and analysis, J.J. and J.N performed bioinformatic analysis, S.M.B,, P.R and A.G planned and / or performed sequencing and analysis, X.Y., J.J., L.R., M.J.G.B. J.V., S.S. and J.P.L wrote the paper

## Methods

### Clinical Samples

Nasal swab specimens from SARS-CoV-2 RT-PCR confirmed cases, selected to be as close as possible to the onset of symptoms and having a SARS-CoV-2 viral load >200 copies/mL, were selected for sequencing. Molecular confirmation of SARS-CoV-2 infection and viral load quantification was performed using the Abbott RealT*ime* SARS-CoV-2 RT-PCR at the Virology Laboratory of the University of Washington, Department of Laboratory Medicine and Pathology (UW Virology, Seattle, US),.

### Next-generation sequencing

Next-generation sequencing (NGS) was performed by UW Virology using the clinically validated Swift Biosciences SNAP Version 2.0 assay (Integrated DNA Technologies). The SNAP assay utilizes multiple overlapping amplicons in a single tube to prepare ready-to-sequence libraries. The primer pairs used in SNAP were designed for generating libraries from first- or second-strand cDNA produced from viral isolates or clinical specimens enabling successful SARS-CoV-2 library preparation from samples with low viral titers. The Swift Biosciences SARS-CoV2 Version 2.0 kit (Catalog # CovG1 V2-96) has been optimized to achieve additional genome coverage on the Illumina sequencing platforms. A full clinical validation with determination of analytical sensitivity and specificity, limit of detection, accuracy, and assay precision (reproducibility and repeatability) has been performed.

### Protein expression and purification

Plasmids corresponding to the SARS-CoV2 S variant proteins truncated after residue 1208 and with stabilizing substitutions A892P, A942P, D614N and V987P and a furin cleavage site knock out (R682S, R685G) were synthesized and codon-optimized at GenScript (Piscataway, NJ 08854). The constructs were cloned into pCDNA2004 or generated by standard methods widely known within the field involving site-directed mutagenesis and PCR and sequenced. The expression platform used was the Expi293F cells. The cells were transiently transfected using ExpiFectamine (Life Technologies) according to the manufacturer’s instructions and cultured for 6 days at 37°C and 10% CO2. The culture supernatant was harvested and spun for 5 minutes at 300 g to remove cells and cellular debris. The spun supernatant was subsequently sterile filtered using a 0.22 μm vacuum filter and stored at 4°C until use. S trimers were purified using a two-step purification protocol by Lentil Lectin from *Galanthus Nivalis* (Vector labs, catalog AL- 1243., followed by by size-exclusion chromatography using a HiLoad Superdex 200 16/600column (GE Healthcare).

### Antibodies and reagents

ACE2-Fc was made according to Liu et al. 2018. Kidney international. For 2-51, DH1055, 4A8, S1M11, S2E12, C144, 2-43 and S309 the heavy and light chain were cloned into a single IgG1 expression vector to express a fully human IgG1 antibody. Antibodies were produced by transfecting the IgG1 expression constructs using the ExpiFectamine™ 293 Transfection Kit (ThermoFisher) in Expi293F (ThermoFisher) cells according to the manufacturer specifications. Purification from serum-free culture supernatants was done using mAb Select SuRe resin (GE Healthcare) followed by rapid desalting using a HiPrep 26/10 Desalting column (GE Healthcare). The final formulation buffer was 20 mM NaAc, 75 mM NaCl, 5% Sucrose pH 5.5. COVA1-22 and COVA2-15 have been kindly provided by Marit van Gils.

### Differential scanning fluorometry (DSF)

0.2 mg of purified protein in 50 μl PBS pH 7.4 (Gibco) was mixed with 15 μl of 20 times diluted SYPRO orange fluorescent dye (5000 x stock, Invitrogen S6650) in a 96-well optical qPCR plate. A negative control sample containing the dye was only used for reference subtraction. The measurement was performed in a qPCR instrument (Applied Biosystems ViiA 7) using a temperature ramp from 25–95 °C with a rate of 0.015 °C per second. Data was collected continuously. The negative first derivative was plotted as a function of temperature. The melting temperature corresponds to the lowest point in the curve.

### BioLayer Interferometry (BLI)

The antibodies were immobilized on anti-hIgG (AHC) sensors (FortéBio cat#18-5060) in 1x kinetics buffer (FortéBio cat#18-1092) in 96-well black flat bottom polypylene microplates (FortéBio cat#3694). The experiment was performed on an Octet RED384 instrument (Pall-FortéBio) at 30□°C with a shaking speed of 1,000□rpm. Activation was 600 s, immobilization of antibodies 900 s, followed by washing for 600 s and then binding the S proteins for 300 s. The data analysis was performed using the FortéBio Data Analysis 12.0 software (FortéBio).

### Cryo-EM Grid Preparation and Data Collection

3.5 μL of 0.8-1.0 mg/ml purified ΔN25 or ΔN135 Spike complex was applied to the plasma-cleaned (Gatan Solarus) Quantifoil 1.2/1.3 holey gold grid, and subsequently vitrified using a Vitrobot Mark IV (FEI Company). Cryo grids were loaded into a Titan Krios transmission electron microscope (ThermoFisher Scientific) with a post-column Gatan Image Filter (GIF) operating in nanoprobe at 300 keV with a Gatan K3 Summit direct electron detector and an energy filter slit width of 20 eV. Images were recorded with Leginon in counting mode with a pixel size of 0.832 Å and a nominal defocus range of −1.8 to −1.2 μm. Images were recorded with a 1.4 s exposure and 40 ms subframes (35 total frames) corresponding to a total dose of ~ 52 electrons per Å2. All details corresponding to individual datasets are summarized in Table S2.

### Cryo-EM image processing

Dose-fractioned movies were gain-corrected, and beam-induced motion correction using MotionCor2(*37*) with the dose-weighting option. The Spike particles were automatically picked from the dose-weighted, motion corrected average images using Relion 3.0(*38*). CTF parameters were determined by Gctf(*39*). Particles were then extracted using Relion 3.0 with a box size of 440 pixels. The 3D classification and refinement were performed with Relion 3.0 using the binned datasets. One round of 3D classification was performed to select the homogenous particles. Unbinned homogenous particles were re-extracted and then submitted to 3D auto-refinement without symmetry imposed. For Brazilian Spike, cryoDRGN was performed using the parameters from the last iteration of the 3D auto-refinement. An additional round of no-alignment 3D classification revealed two distinct conformational states of ΔN135 Spike: ~73 % of particles adopting an open conformation with one erected RBD was further refined without symmetry imposed; ~23 % of particles in the fully closed conformation were further refined with the C3 symmetry imposed. An additional round of no-alignment 3D classification revealed one open state of ΔN25 Spike and was followed by further refinement without symmetry imposed. Focus refinements were performed with soft masks around the NTD, RBD, and body regions. 3D classifications and 3D refinements were started from a 60 Å low-pass filtered version of an ab initio map generated with Relion 3.0. All resolutions were estimated by applying a soft mask around the protein complex density and based on the gold-standard (two halves of data refined independently) FSC□=□0.143 criterion. Prior to visualization, all density maps were sharpened by applying different negative temperature factors using automated procedures, along with the half maps, were used for model building. Local resolution was determined using ResMap(*40*) (Fig. S5).

### Model building and refinement

The initial template of the Spike complex was derived from a homology-based model calculated by SWISS-MODEL(*41*). The model was docked into the EM density map using Chimera(*42*) and followed by manually adjustment using COOT(*43*). Note that the EM density around the NTD and RBD regions was poor relative to other parts of the model. The NTD and RBD regions were modeled using the unsharpened maps together with the deepEMhancer maps that were calculated with the half maps from the focus refinements. Each model was independently subjected to global refinement and minimization in real space using the module phenix.real_space_refine in PHENIX(*44*) against separate EM half-maps with default parameters. The model was refined into a working half-map, and improvement of the model was monitored using the free half map. Model geometry was further improved using Rosetta. The geometry parameters of the final models were validated in COOT and using MolProbity(*45*)and EMRinger(*46*). These refinements were performed iteratively until no further improvements were observed. The final refinement statistics were provided in Table S2. Model overfitting was evaluated through its refinement against one cryo-EM half map. FSC curves were calculated between the resulting model and the working half map as well as between the resulting model and the free half and full maps for cross-validation (Figure S6). Figures were produced using PyMOL (The PyMOL Molecular Graphics System) and Chimera.

### Analytical SEC

An ultra-high-performance liquid chromatography system (Vanquish, Thermo Scientific) and μDAWN TREOS instrument (Wyatt) coupled to an Optilab μT-rEX Refractive Index Detector (Wyatt), in combination with an in-line Nanostar DLS reader (Wyatt), was used for performing the analytical SEC experiment. The cleared crude cell culture supernatants were applied to a SRT-10C SEC-500 15 cm column, (Sepax Cat# 235500-4615) with the corresponding guard column (Sepax) equilibrated in running buffer (150 mM sodium phosphate, 50 mM NaCl, pH 7.0) at 0.35 mL/min. When analyzing supernatant samples, μMALS detectors were offline and analytical SEC data was analyzed using Chromeleon 7.2.8.0 software package. The signal of supernatants of non-transfected cells was subtracted from the signal of supernatants of S transfected cells. When purified proteins were analyzed using SEC-MALS, μMALS detectors were inline and data was analyzed using Astra 7.3 software package.

### Cell-cell fusion assay

A GFP-based cell-cell fusion assay was performed to determine the capability of the variant S protein to mediate membrane fusion. HEK293 cells were transfected with full-length S, human ACE2, human TMPRSS2 and GFP. All proteins were expressed from pcDNA2004 plasmids using Trans-IT transfection reagent according to the manufacturer’s instructions. 18hr after transfection, syncytia formation was visualized on an EVOS microscope.

### GISAID data acquisition and processing

SARS-CoV-2 genome and sample data were downloaded from the GISAID Initiative (https://www.gisaid.org/) database on 25 Jan 2022, and processed by Biovia Pipeline Pilot workflows (BIOVIA, Dassault Systèmes, v 21.2.0.2574, San Diego: Dassault Systèmes, 2020) to transform and standardize the date and country formats, and to retain only human samples. The data are subsequently saved to files with information on individual lineages and individual mutations in Spike protein The data was further analyzed in Tableau (www.tableau.com) to obtain mutation and lineage frequencies as function of time or location.

Phylogenetic trees in Fig 6A and Fig 6C were created using amino-acid sequences of the S-proteins from GISAID. For each lineage, only one, the most frequent S-protein sequence was used. Only lineages that had 50 or more identical sequences store od GISAID as of 25 Jan 2022 were used. The trees were created using the CLC software.

## References

1. W. H. Chen, P. J. Hotez, M. E. Bottazzi, Potential for developing a SARS-CoV receptor-binding domain (RBD) recombinant protein as a heterologous human vaccine against coronavirus infectious disease (C0VID)-19. Hum Vaccin Immunother, 1–4 (2020).

2. L. Liu et al., Potent neutralizing antibodies against multiple epitopes on SARS-CoV-2 spike. Nature 584, 450–456 (2020).

3. M. Yuan et al., A highly conserved cryptic epitope in the receptor binding domains of SARS-CoV-2 and SARS-CoV. Science 368, 630–633 (2020).

4. P. J. M. Brouwer et al., Potent neutralizing antibodies from COVID-19 patients define multiple targets of vulnerability. Science, (2020).

5. B. J. Bosch, R. van der Zee, C. A. de Haan, P. J. Rottier, The coronavirus spike protein is a class I virus fusion protein: structural and functional characterization of the fusion core complex. J Virol 77, 8801–8811 (2003).

6. F. Li, Structure, Function, and Evolution of Coronavirus Spike Proteins. Annu Rev Virol 3, 237–261 (2016).

7. D. Wrapp et al., Cryo-EM structure of the 2019-nCoV spike in the prefusion conformation. Science 367, 1260–1263 (2020).

8. K. M. Hastie et al., Defining variant-resistant epitopes targeted by SARS-CoV-2 antibodies: A global consortium study. Science 374, 472–478 (2021).

9. W. N. Voss et al., Prevalent, protective, and convergent IgG recognition of SARS-CoV-2 non-RBD spike epitopes. Science 372, 1108–1112 (2021).

10. X. Chi et al., A neutralizing human antibody binds to the N-terminal domain of the Spike protein of SARS-CoV-2. Science 369, 650–655 (2020).

11. D. Li et al., In vitro and in vivo functions of SARS-CoV-2 infection-enhancing and neutralizing antibodies. Cell 184, 4203–4219 e4232 (2021).

12. N. Suryadevara et al., Neutralizing and protective human monoclonal antibodies recognizing the N-terminal domain of the SARS-CoV-2 spike protein. Cell 184, 2316–2331 e2315 (2021).

13. M. McCallum et al., N-terminal domain antigenic mapping reveals a site of vulnerability for SARS-CoV-2. Cell 184, 2332–2347 e2316 (2021).

14. G. Cerutti et al., Potent SARS-CoV-2 neutralizing antibodies directed against spike N-terminal domain target a single supersite. Cell Host Microbe 29, 819–833 e817 (2021).

15. D. Haslwanter et al., A Combination of Receptor-Binding Domain and N-Terminal Domain Neutralizing Antibodies Limits the Generation of SARS-CoV-2 Spike Neutralization-Escape Mutants. mBio 12, e0247321 (2021).

16. M. L. Acevedo et al., Infectivity and immune escape of the new SARS-CoV-2 variant of interest Lambda. medRxiv, 2021.2006.2028.21259673 (2021).

17. M. Hoffmann et al., SARS-CoV-2 variants B.1.351 and P.1 escape from neutralizing antibodies. Cell 184, 2384–2393 e2312 (2021).

18. M. McCallum et al., Molecular basis of immune evasion by the Delta and Kappa SARS-CoV-2 variants. Science 374, 1621–1626 (2021).

19. D. Mannar et al., SARS-CoV-2 Omicron variant: Antibody evasion and cryo-EM structure of spike protein-ACE2 complex. Science, eabn7760 (2022).

20. J. Sadoff et al., Final Analysis of Efficacy and Safety of Single-Dose Ad26.COV2.S. N Engl J Med, (2022).

21. R. Bos et al., Ad26 vector-based COVID-19 vaccine encoding a prefusion-stabilized SARS-CoV-2 Spike immunogen induces potent humoral and cellular immune responses. NPJ Vaccines 5, 91 (2020).

22. J. Juraszek et al., Stabilizing the closed SARS-CoV-2 spike trimer. Nat Commun 12, 244 (2021).

23. M. A. Tortorici et al., Ultrapotent human antibodies protect against SARS-CoV-2 challenge via multiple mechanisms. Science 370, 950–957 (2020).

24. C. O. Barnes et al., SARS-CoV-2 neutralizing antibody structures inform therapeutic strategies. Nature 588, 682–687 (2020).

25. D. Pinto et al., Cross-neutralization of SARS-CoV-2 by a human monoclonal SARS-CoV antibody. Nature 583, 290–295 (2020).

26. P. J. M. Brouwer et al., Potent neutralizing antibodies from COVID-19 patients define multiple targets of vulnerability. Science 369, 643–650 (2020).

27. R. E. Chen et al., Resistance of SARS-CoV-2 variants to neutralization by monoclonal and serum-derived polyclonal antibodies. Nat Med 27, 717–726 (2021).

28. S. Iketani et al., Lead compounds for the development of SARS-CoV-2 3CL protease inhibitors. Nat Commun 12, 2016 (2021).

29. M. Yuan et al., Structural and functional ramifications of antigenic drift in recent SARS-CoV-2 variants. Science 373, 818–823 (2021).

30. I. Kimura et al., The SARS-CoV-2 Lambda variant exhibits enhanced infectivity and immune resistance. Cell Rep 38, 110218 (2022).

31. S. M. Gobeil et al., Effect of natural mutations of SARS-CoV-2 on spike structure, conformation, and antigenicity. Science 373, (2021).

32. M. McCallum et al., SARS-CoV-2 immune evasion by the B.1.427/B.1.429 variant of concern. Science 373, 648–654 (2021).

33. H. Nielsen, K. D. Tsirigos, S. Brunak, G. von Heijne, A Brief History of Protein Sorting Prediction. Protein J 38, 200–216 (2019).

34. https://www.ecdc.europa.eu/en/covid-19/variants-concern.

35. C. Scheepers et al., Emergence and phenotypic characterization of C.1.2, a globally detected lineage that rapidly accumulated mutations of concern. medRxiv, 2021.2008.2020.21262342 (2021).

36. P. Colson et al., Emergence in Southern France of a new SARS-CoV-2 variant of probably Cameroonian origin harbouring both substitutions N501Y and E484K in the spike protein. medRxiv, 2021.2012.2024.21268174 (2021).

37. S. Q. Zheng et al., MotionCor2: anisotropic correction of beam-induced motion for improved cryo-electron microscopy. Nat Methods 14, 331–332 (2017).

38. J. Zivanov et al., New tools for automated high-resolution cryo-EM structure determination in RELI0N-3. Elife 7, (2018).

39. K. Zhang, Gctf: Real-time CTF determination and correction. J Struct Biol 193, 1–12 (2016).

40. A. Kucukelbir, F. J. Sigworth, H. D. Tagare, Quantifying the local resolution of cryo-EM density maps. Nat Methods 11, 63–65 (2014).

41. A. Waterhouse et al., SWISS-MODEL: homology modelling of protein structures and complexes. Nucleic Acids Res 46, W296–W303 (2018).

42. E. F. Pettersen et al., UCSF Chimera--a visualization system for exploratory research and analysis. J Comput Chem 25, 1605–1612 (2004).

43. P. Emsley, B. Lohkamp, W. G. Scott, K. Cowtan, Features and development of Coot. Acta Crystallogr D Biol Crystallogr 66, 486–501 (2010).

44. P. V. Afonine et al., Real-space refinement in PHENIX for cryo-EM and crystallography. Acta Crystallogr D Struct Biol 74, 531–544 (2018).

45. V. B. Chen et al., MolProbity: all-atom structure validation for macromolecular crystallography. Acta Crystallogr D Biol Crystallogr 66, 12–21 (2010).

46. B. A. Barad et al., EMRinger: side chain-directed model and map validation for 3D cryo-electron microscopy. Nat Methods 12, 943–946 (2015).

