## Supplemental Files for "Convergence of immune escape strategies highlights plasticity of SARS-CoV-2 spike"

**Table S1. Overview of collected sequences with large deletions in the spike variants from the COV3001 trial**

| Sequence ID | Collection date | Country code | Lineage | #Deletions | #Mutations | P9L | S46L | **T63del** | **Del 64-75** | **T76del** | **Del 136-144** | **R246del** | R246N | **Del 247-252** | D253N | **D253del** | **Del 258-264** | D287N | L452Q | E471Q | E484K | F490S | S494P | D614G | N703S | T859N | V1176F |
| --- | --- | --- | --- | --- | --- | --- | --- | --- | --- | --- | --- | --- | --- | --- | --- | --- | --- | --- | --- | --- | --- | --- | --- | --- | --- | --- | --- |
| 1 | 12-Jan-21 | BR | B.1.1.294 | 16 | 21 | x |  |  |  |  | ● |  |  |  |  |  | ● |  |  |  | x |  | x | x |  |  | x |
| 2 | 13-Jan-21 | PE | C.37 | 20 | 26 |  |  |  | ● | ● |  |  | x | ● |  | ● |  |  | x | x |  | x |  | x |  | x |  |
| 3 | 17-Jan-21 | BR | B.1.1.294 | 16 | 21 | x |  |  |  |  | ● |  |  |  |  |  | ● |  |  |  | x |  | x | x |  |  | x |
| 4 | 9-Feb-21 | PE | C.37 | 20 | 25 |  |  |  | ● | ● |  |  | x | ● |  | ● |  |  | x |  |  | x |  | x |  | x |  |
| 5 | 23-Feb-21 | PE | C.37 | 20 | 26 |  |  | ● | ● |  |  | ● |  | ● | x |  |  |  | x | x |  | x |  | x |  | x |  |
| 6 | 24-Feb-21 | PE | C.37 | 20 | 26 |  |  | ● | ● |  |  | ● |  | ● | x |  |  |  | x | x |  | x |  | x |  | x |  |
| 7 | 16-Mar-21 | PE | C.37 | 20 | 25 |  |  | ● | ● |  |  | ● |  | ● | x |  |  |  | x |  |  | x |  | x |  | x |  |
| 8 | 2-Apr-21 | PE | C.37 | 20 | 27 |  | x | ● | ● |  |  | ● |  | ● | x |  |  |  | x |  |  | x |  | x | x | x |  |
| 9 | 5-Apr-21 | AR | C.37 | 20 | 26 |  |  | ● | ● |  |  | ● |  | ● | x |  |  | x | x |  |  | x |  | x |  | x |  |
| 10 | 15-Apr-21 | PE | C.37 | 20 | 25 |  |  | ● | ● |  |  | ● |  | ● | x |  |  |  | x |  |  | x |  | x |  | x |  |
| 11 | 26-May-21 | PE | C.37 | 20 | 25 |  |  | ● | ● |  |  | ● |  | ● | x |  |  |  | x |  |  | x |  | x |  | x |  |
| 12 | 25-Jun-21 | AR | C.37 | 20 | 25 |  |  | ● | ● |  |  | ● |  | ● | x |  |  |  | x |  |  | x |  | x |  | x |  |

**Table S2. Data collection, reconstruction, and model refinement statistics**

|  | ΔN135 (C1)  EMD-xxxx, PDB: xxxx | ΔN135 (C3)  EMD-xxxx, PDB: xxxx | ΔN25 (C1)  EMD-xxxx, PDB: xxxx |
| --- | --- | --- | --- |
| **Data collection** |  |  |  |
| Microscope | Titan Krios | | Titan Krios |
| Voltage (keV) | 300 | | 300 |
| Nominal magnification | 105000 x | | 105000 x |
| Exposure navigation | Image Shift | | Image Shift |
| Electron exposure (e /Å^2^) | 51.98 | | 52.25 |
| Total exposure time (sec) | 1.4 | | 1.4 |
| Detector | K3 Summit | | K3 Summit |
| Pixel size (Å)* | 0.832 | | 0.832 |
| Defocus range (µm) | -1.2 to -1.8 | | -1.2 to -1.6 |
| Micrographs Used | 20,732 | | 18,021 |
| Final Refined particles (no.) | 378,064 | 115,817 | 207,525 |
| **Reconstruction** |  |  |  |
| Symmetry imposed | C1 | C3 | C1 |
| Resolution (global) |  |  |  |
| FSC 0.143 | 3.08 Å | 3.21 Å | 3.52 Å |
| Applied B-factor (Å^2^) | -101 | -118 | -125 |
| **Refinement** |  |  |  |
| Protein residues | 3137 | 3108 | 3147 |
| Map Correlation Coefficient (Main chain) | 0.8439 | 0.8216 | 0.7131 |
| R.m.s deviations |  |  |  |
| Bond lengths (Å) | 0.012 | 0.012 | 0.011 |
| Bond angles (°) | 1.08 | 1.08 | 1.09 |
| Ramachandran |  |  |  |
| Outliers | 1.05 % | 0.87 % | 0.76 % |
| Allowed | 5.36 % | 5.18 % | 5.63 % |
| Favored | 93.59 % | 93.95 % | 93.61 % |
| Rotamer outliers | 0.07 % | 0.11 % | 0.07 % |
| MolProbity score | 1.36 | 1.35 | 1.36 |
| EMRinger score | 3.35 | 2.82 | 1.94 |
| Clashscore (all atoms) | 1.77 | 1.84 | 1.77 |

*Calibrated pixel size at the detector

**Table S3. Mutations leading to DS_15-136_ loss in lineages**

| **Mutation SP** | **Mutation 136** | **Count** | **Lineage** | **Percent of mutation within lineage** |
| --- | --- | --- | --- | --- |
| S13I | WT | 38,910 | B.1.429 | 82.33% |
| S13I | WT | 16,514 | B.1.427 | 73.51% |
| S13I | WT | 877 | B.1.429 | 1.86% |
| S13I | WT | 736 | B.1.427 | 3.28% |
| L5F+S13I | WT | 669 | B.1.429 | 1.42% |
| L5F+S13I | WT | 468 | B.1.427 | 2.08% |
| P9L+S13I | WT | 310 | B.1.427 | 1.38% |
| P9L | C136del | 267 | B.1.640.1 | 56.21% |
| P9L | C136F | 222 | C.1.2 | 83.15% |
| P9L | WT | 203 | B.1.640.1 | 42.74% |
| P9L | C136F | 171 | B.1.630 | 87.24% |
| P9L | WT | 167 | P.3 | 27.24% |
| P9L | C136del | 140 | AT.1 | 83.33% |
| P9L | WT | 103 | C.38 | 66.88% |
| S13I | WT | 43 | A.2.5 | 1.99% |
| S13I | WT | 35 | B.1.429.1 | 64.81% |
| P9L | WT | 30 | N.10 | 88.24% |
| P9L | WT | 22 | C.1.2 | 8.24% |
| WT | C136F | 20 | C.1.2 | 7.49% |
| P9L | WT | 20 | C.39 | 39.22% |
| P9L | WT | 19 | B.1.1.524 | 38.78% |
| S12F+S13I | WT | 19 | C.36.3 | 1.05% |
| P9L | C136F | 18 | B.1.1.524 | 36.73% |
| P9L | WT | 17 | B.1.640.2 | 48.57% |
| P9L | C136F | 12 | B.1.639 | 9.92% |
| P9L | WT | 12 | P.3 | 1.96% |
| P9L | C136del | 11 | B.1.640.2 | 31.43% |
| P9L | WT | 10 | AT.1 | 5.95% |
| P9L | WT | 8 | B.1.630 | 4.08% |
| WT | C136F | 7 | B.1.630 | 3.57% |
| L5F+Q14del+C15del | C136Y | 7 | B.1.638 | 6.93% |
| L5F+P9L | C136del | 7 | B.1.640.2 | 20.00% |
| P9L | C136F | 5 | B.1.1.10 | 1.16% |
| P9L | WT | 4 | B.1.149 | 1.63% |
| WT | C136F | 4 | B.1.639 | 3.31% |
| P9L | WT | 4 | C.38 | 2.60% |
| L8F+P9L | WT | 3 | AZ.2.1 | 2.36% |
| C15S | C136F | 3 | B.1.1.524 | 6.12% |
| P9L | WT | 3 | B.1.482 | 50.00% |
| Q14del+C15del | WT | 3 | W.1 | 3.09% |

**Table S4. Local occurrences of DS_15-136_ loss in the Delta and Omicron lineages.**

| **Lineage** | **C15*** | **C136** | **both** | **Country or state** | **Period** | **#sequences** | **Lineage percentage** |
| --- | --- | --- | --- | --- | --- | --- | --- |
| BA.1.1 |  | x |  | US-Idaho | Q1 2022 | 249 | 9.6% |
| AY.127 | x |  |  | Sweden | Q3 2021 | 8 | 6.6% |
| AY.34.1.1 |  | x |  | Chile | Q3 2021 | 5 | 5.1% |
| BA.1 | x |  |  | US-Alaska | Q1 2022 | 11 | 3.7% |
| BA.1.1 |  | x |  | US-Louisiana | Q1 2022 | 18 | 2.0% |
| BA.1 |  | x |  | Greece | Q4 2021 | 5 | 1.9% |
| BA.1.1 | x | x |  | US-California | Q1 2022 | 370 | 1.3% |

**Supplementary Figure 1. Differential scanning fluorimetry.** Analysis of melting temperature (Tm) using differential scanning fluorimetry of purified S protein Wuhan-Hu-1 (A), ΔN135 (B) and ΔN25 (C) variants. The first order derivatives are plotted. The experiment was done in triplicate. The Tm is determined as the lowest derivative value representing the Tm50 value

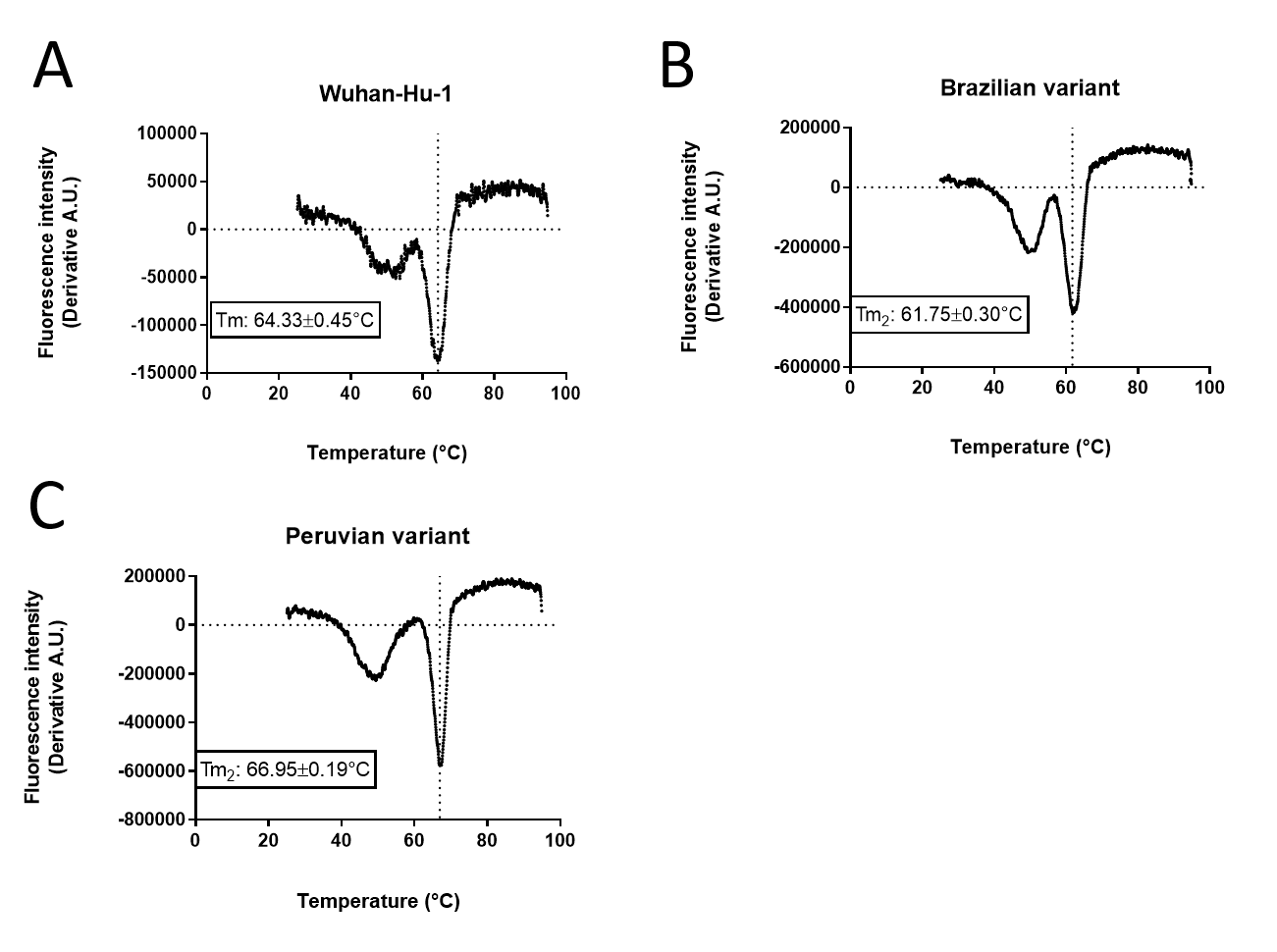

ΔN25

ΔN135

**Supplementary Figure 2.** Evolutionary relationships among SARS1-like coronaviruses (blue) and SARS-CoV-2-like coronaviruses (red) with the N-terminal residues of spike of the indicated branch in green

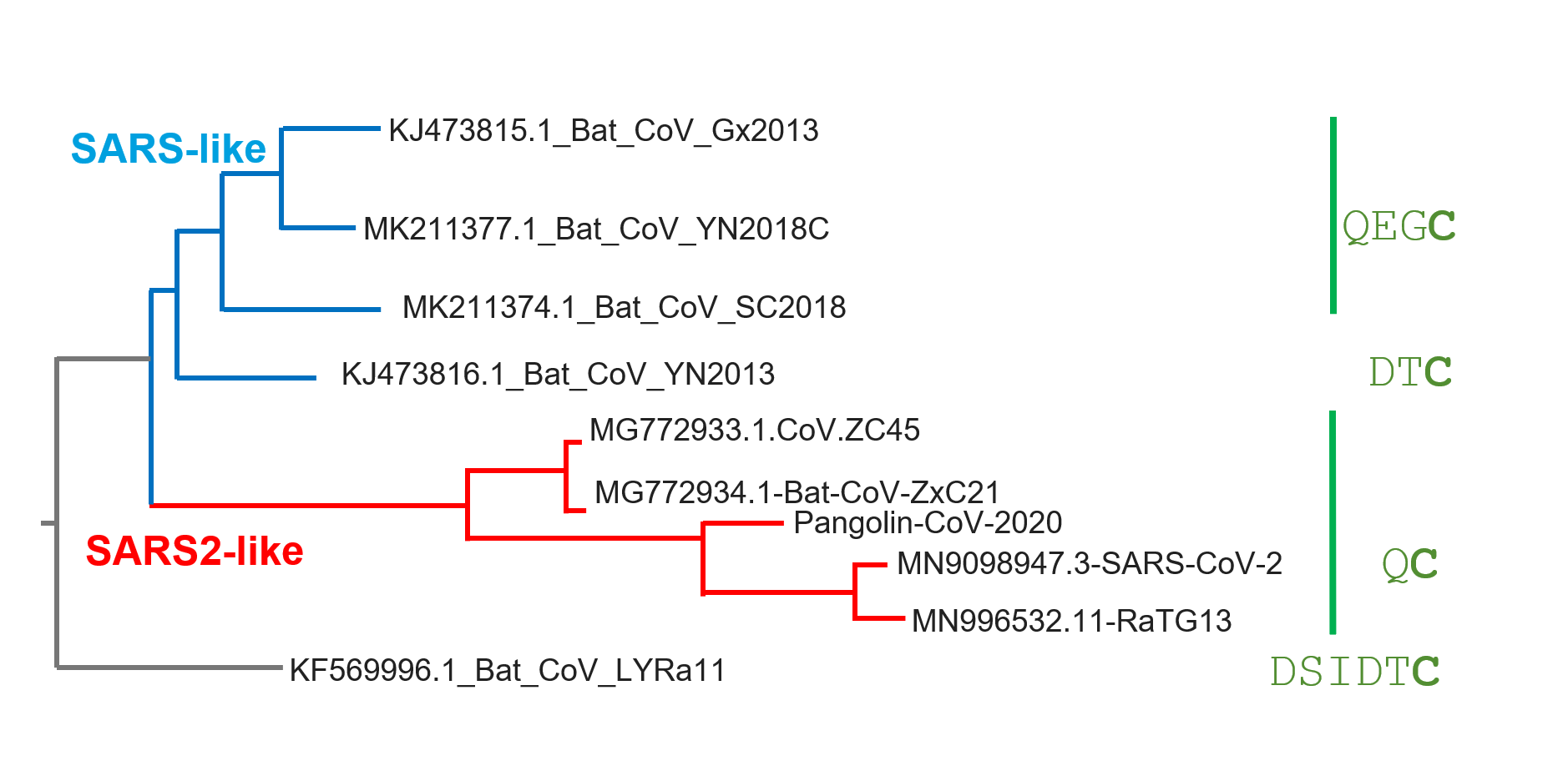

**Supplementary Figure 3.** Mass spectrometry (MS) analysis of selected peptides. A-B. ESI-MS (A) and MS/MS (B) spectrum of the most N-terminal peptide observed of wild type S-protein treated with trypsin protease. MS/MS analysis shows the MS plot with the most prominent peaks labelled (left) and a list of the identified fragmented peptides (right). C-D. ESI-MS (C) and MS/MS (D) spectrum of the most N-terminal peptide observed of Brazilian variant S-protein treated with trypsin protease. MS/MS analysis shows the MS plot with the most prominent peaks labelled (left) and a list of the identified fragmented peptides (right).

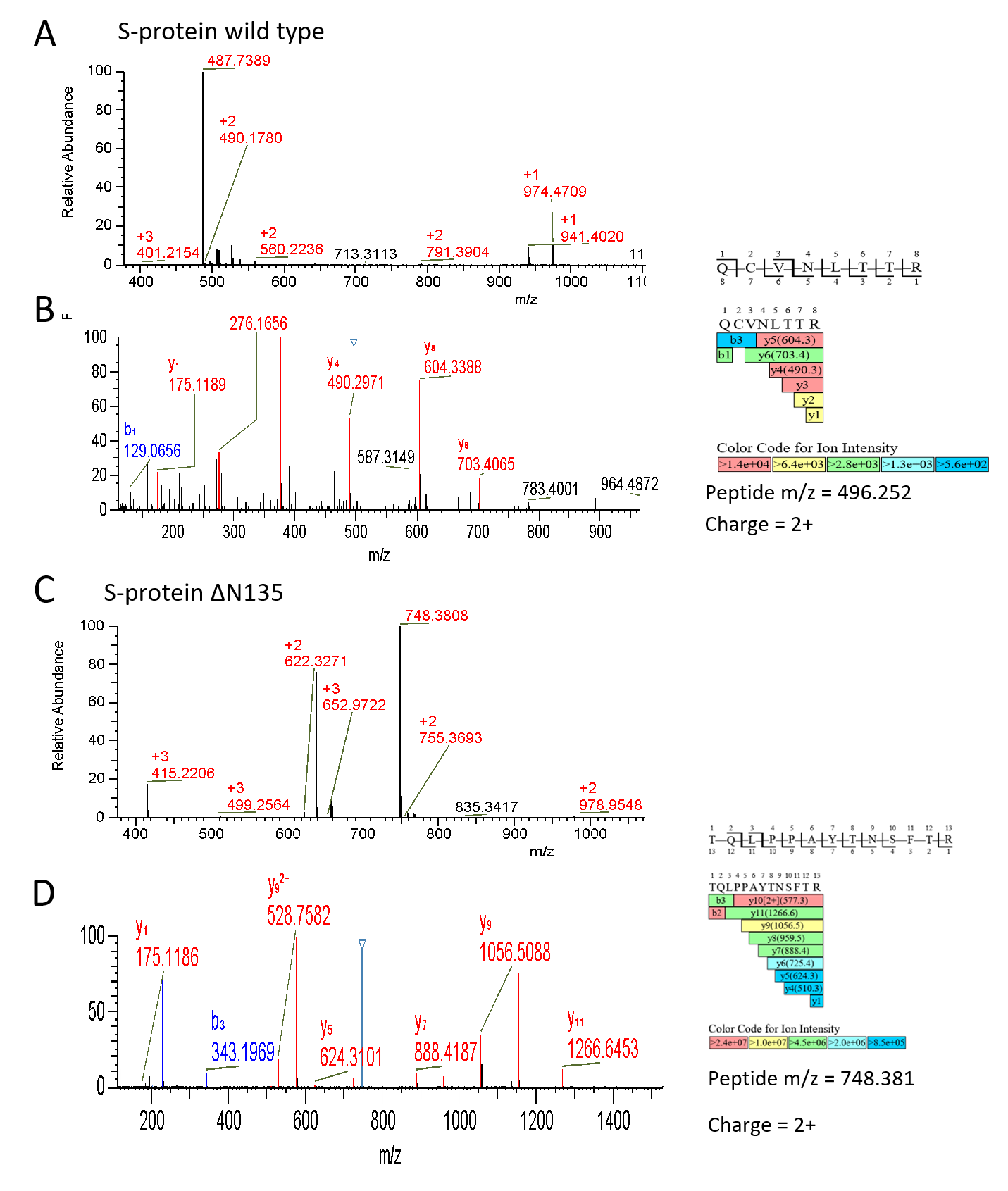

**Supplementary Figure 4.** Cryo-EM structures of ΔN135 and ΔN25 Spikes. (A) Schematic of 2019-nCoV S primary structure colored by domain. Locations of the deletions were highlighted. (B) Density maps of Spike homotrimer viewed from the side highlighting the NTD (navy), RBD (navy), SD1/2 (green) and N-acetyl-D-glucosamine (NAG, in orange). deepEMhancer map focusing on the erect RBD of ΔN25 spike was shown.

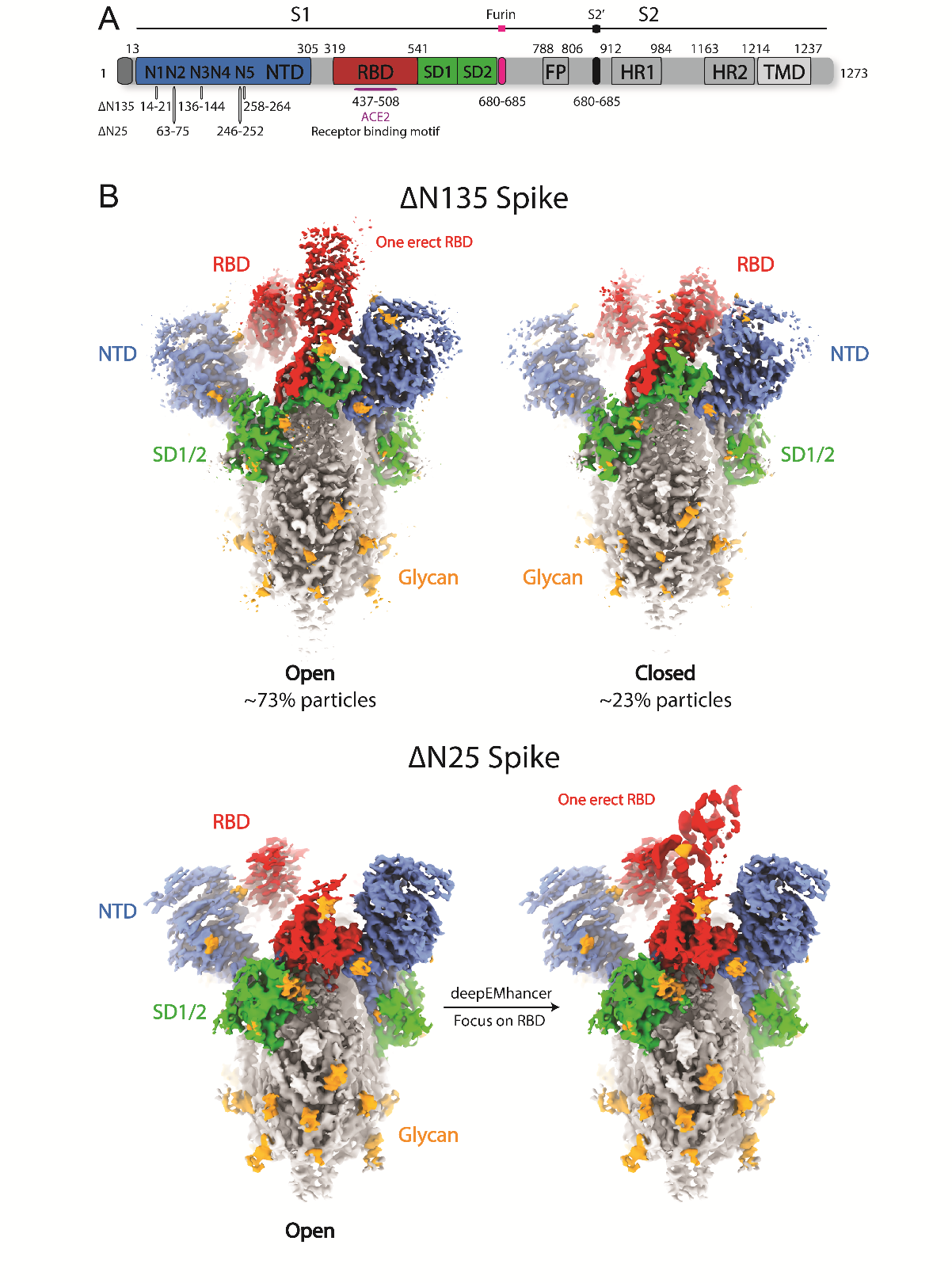

**Supplementary Figure 5a.** Cryo-EM analysis of ΔN25 Spike complex. (A) Flow chart of the cryo-EM data processing procedure. Details can be found in the Materials and methods. (B) A representative cryo-EM micrograph. (C) Angular orientation distribution of the particles used in the final reconstruction. The particle distribution is indicated by different color shades. (D) Local resolution of the map estimated using the ResMap program and colored as indicated. (E) Fourier shell correlation (FSC) curve of the structure with FSC as a function of resolution using Relion output. The resolution is ~3.52 Å at the FSC cutoff of 0.143.

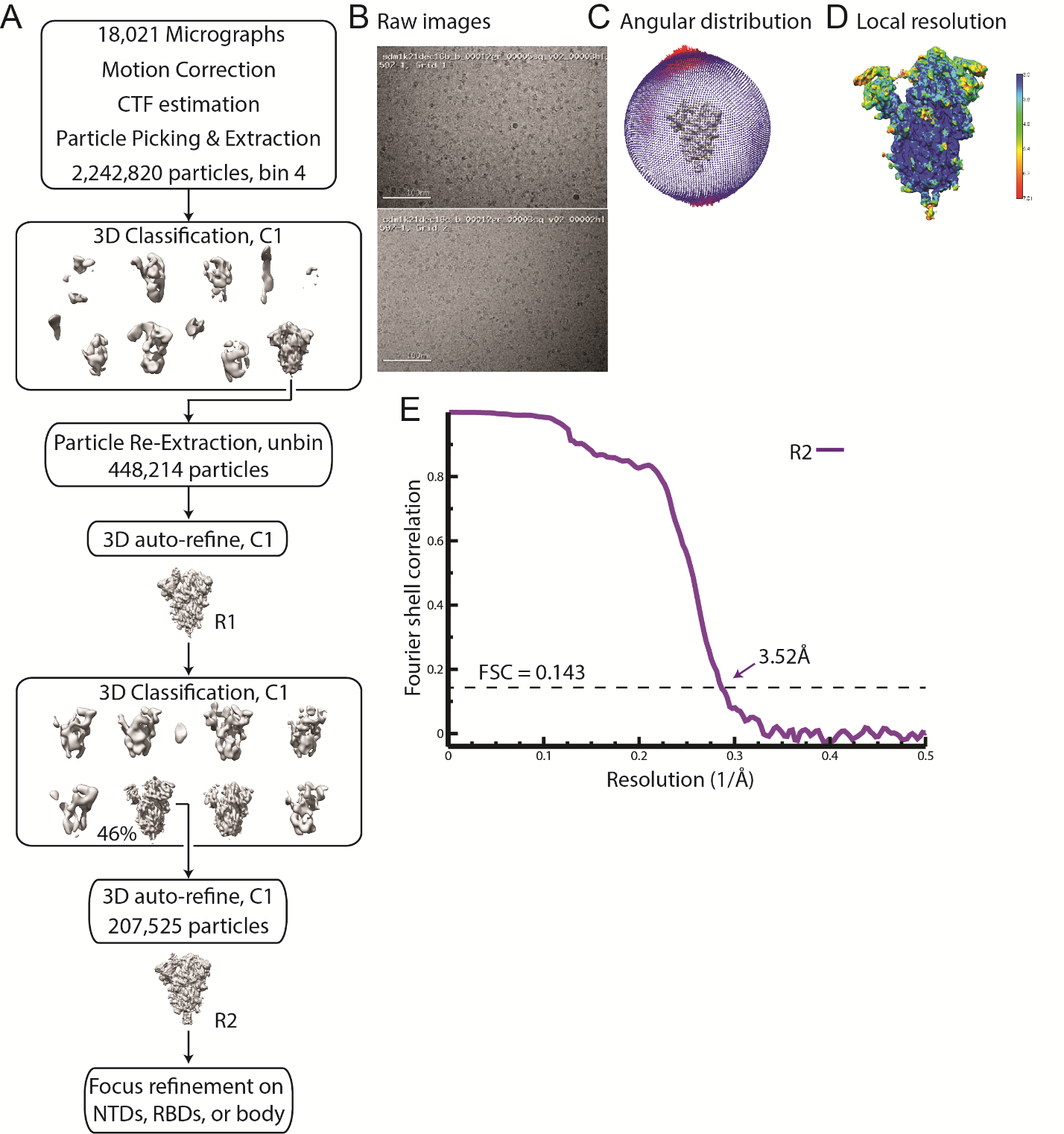

**Supplementary Figure 5b.** Cryo-EM analysis of ΔN135 Spike complex. (A) Flow chart of the cryo-EM data processing procedure. Details can be found in the Materials and methods. (B) A representative cryo-EM micrograph. (C) Angular orientation distribution of the particles used in the final reconstruction. The particle distribution is indicated by different color shades. (D) Local resolution of the map estimated using the ResMap program and colored as indicated. (E) Fourier shell correlation (FSC) curve of the structure with FSC as a function of resolution using Relion output. The resolutions are ~3.08 Å and 3.21 Å at the FSC cutoff of 0.143 for the RBD 1-up and all closed Spikes, respectively. (F) cryoDRGN analysis of the ΔN135 spike revealing the mobility of the RBD highlighted in red arrow.

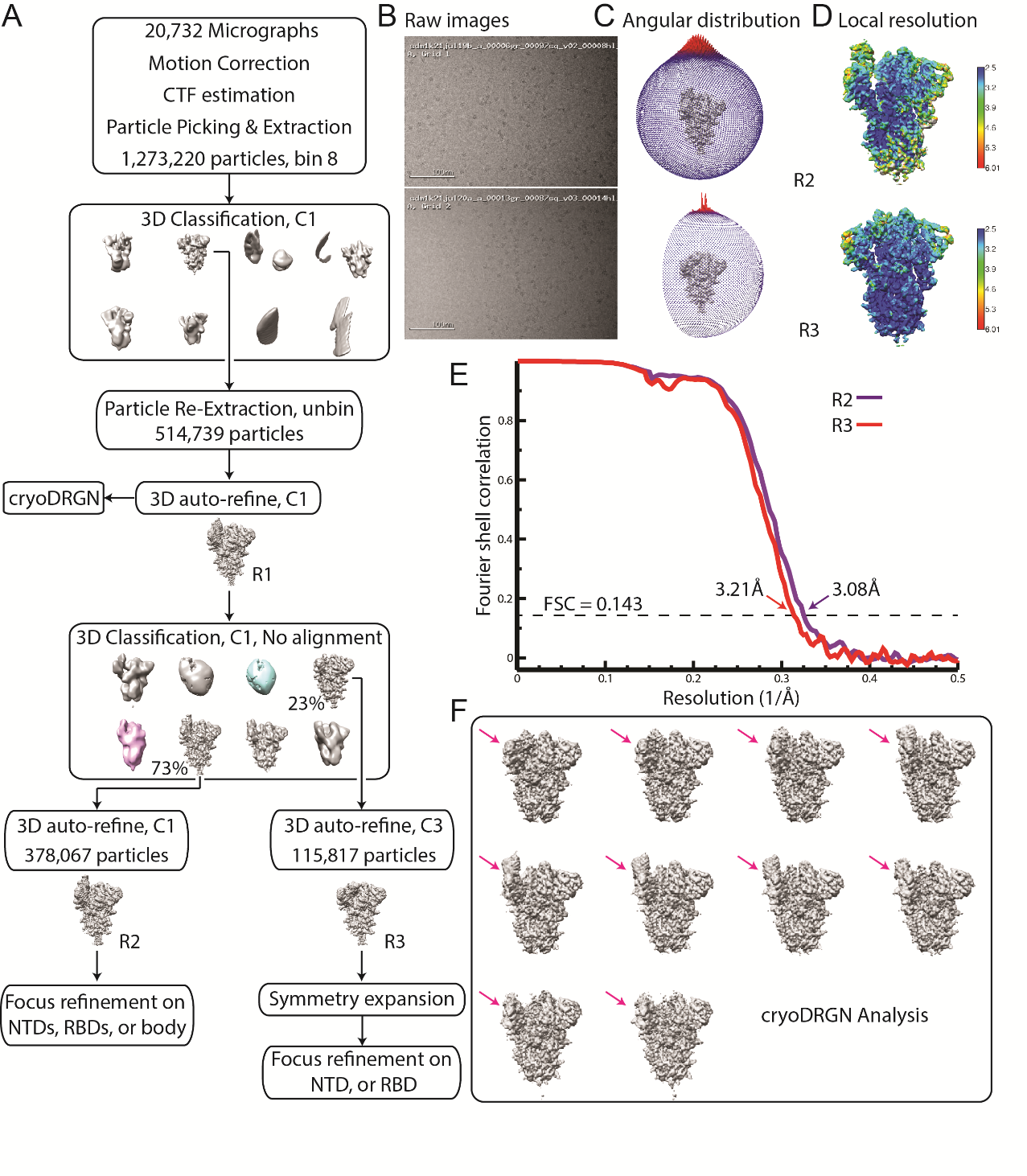

**Supplementary Figure 6.** Cryo-EM densities of the Spike complex. (A) Model validation. Comparison of the FSC curves between model and half map 1 (work), model and half map 2 (free), and model and full map are plotted in red, green and blue, respectively. (B) Representative sharpened Cryo-EM density is displayed as mesh at the contour level 15σ. The atomic model with side chains is shown as sticks. (C) Representative unsharpened Global refinement, Focus Refinement, and deepEMhancer maps around NTD and RBD were shown as surface at 4.5 σ, respectively.

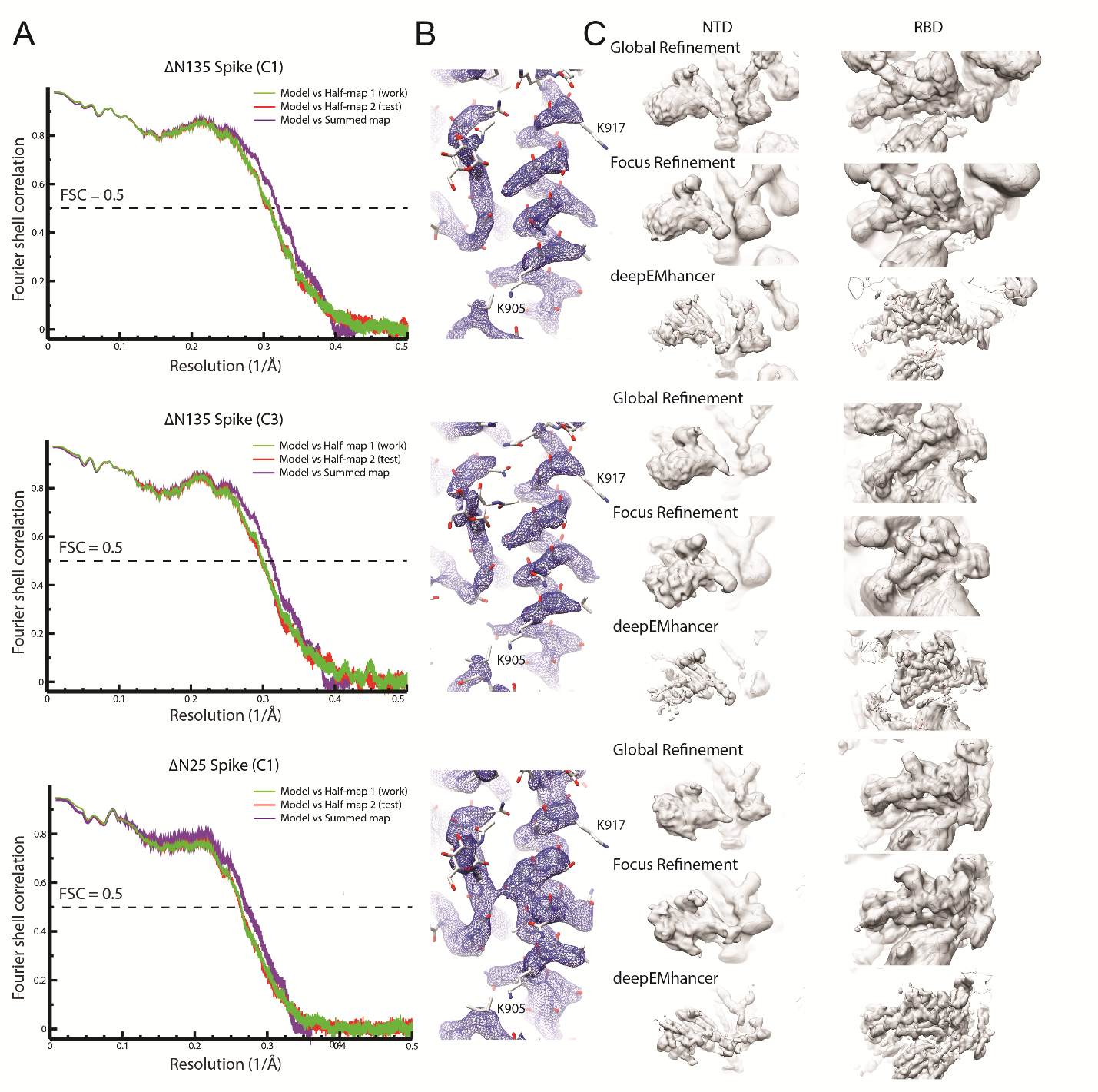

**Supplementary Figure 7.** Superposition of wire model representation of ΔN135 (top) and ΔN25 (bottom) NTD compared with Wuhan (WT) and other lineages with colors and view as in Fig 1B. Arrows indicate rearrangement of the loops as a result of the deletions.

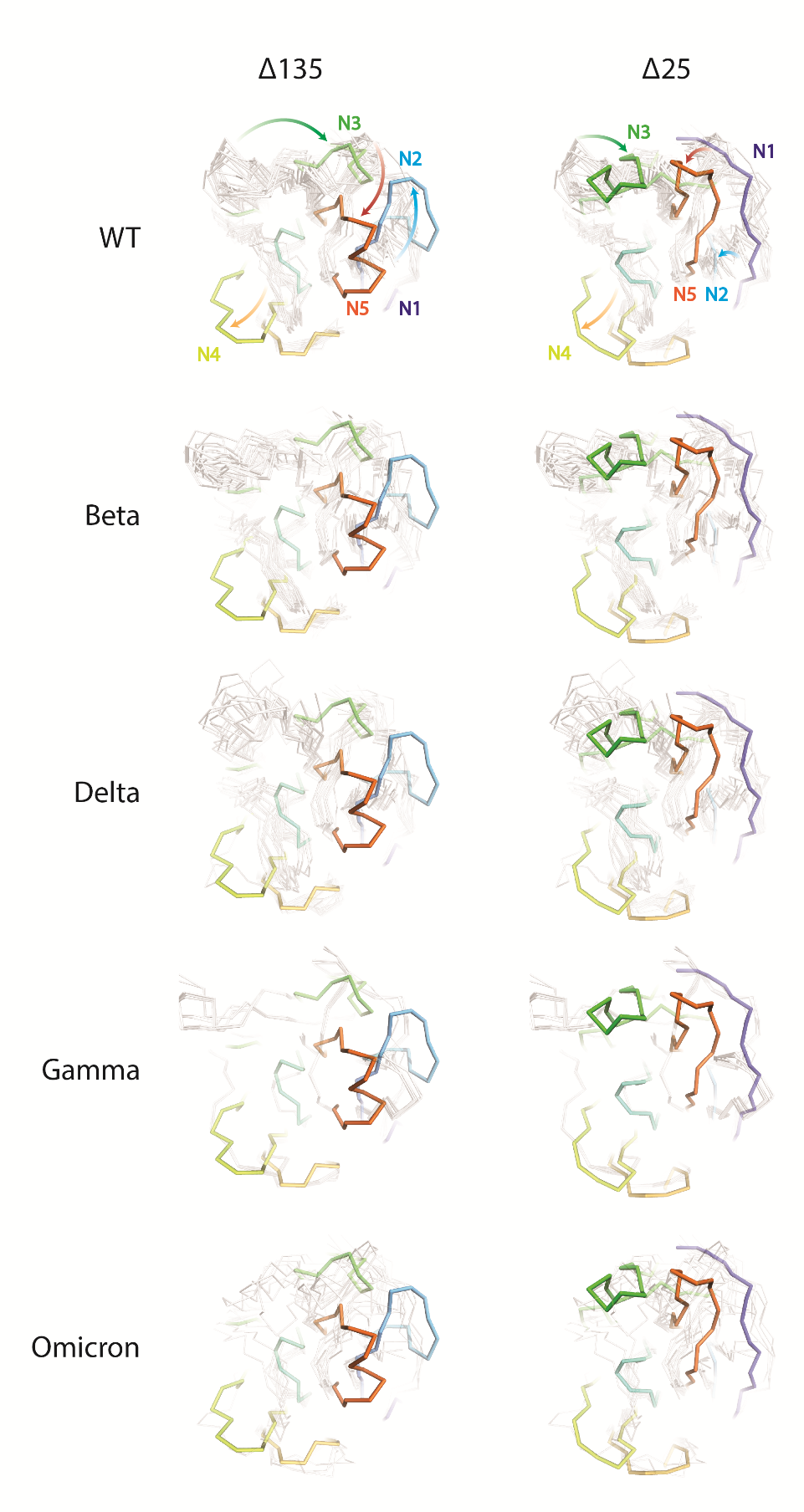
